# Detection of selection signatures in farmed coho salmon (*Oncorhynchus kisutch*) using dense genome-wide information

**DOI:** 10.1101/2020.07.22.215988

**Authors:** M.E. López, M.I. Cádiz, E.B. Rondeau, B.F. Koop, J.M. Yáñez

**Affiliations:** Department of Aquatic Resources, Swedish University of Agricultural Sciences, Drottningholm, Sweden; Facultad de Ciencias Veterinarias y Pecuarias, Universidad de Chile, Santiago, Chile; Department of Biology, University of Victoria, British Columbia, Canada; Núcleo Milenio INVASAL, Concepción, Chile

## Abstract

Animal domestication and artificial selection give rise to gradual changes at the genomic level in populations. Subsequent footprints of selection known as selection signatures or selective sweeps have been traced in the genomes of many animal livestock species by exploiting variations in linkage disequilibrium patterns and/or reduction of genetic diversity.

Domestication of most aquatic species is recent in comparison with land animals, and salmonids are one of the most important fish species in aquaculture. Coho salmon (*Oncorhynchus kisutch*), cultivated primarily in Chile, has been subject to breeding programs to improve growth, disease resistance traits, and flesh color. This study aimed to identify selection signatures that may be involved in adaptation to culture conditions and traits of productive interest. To do so, individuals of two domestic populations cultured in Chile were genotyped with 200 thousand SNPs, and analyses were conducted using iHS, XP-EHH and CLR. Several signatures of selection on different chromosomal regions were detected across both populations. Some of the identified regions under selection contained genes such *anapc2*, *alad*, *chp2* and *myn* that have been previously associated with body weight in Atlantic salmon or *sec24d* and *robo1* that have been associated with disease resistance to *Piscirickettsia salmonis* in coho salmon. Findings in our study can contribute to an integrated genome-wide map of selection signatures, to help identify the genetic mechanisms of phenotypic diversity in coho salmon.

## Introduction

The implementation of genetic improvement programs in domestic populations to improve desirable production traits, combined with the natural adaptation to new environments, has led to a variety of phenotypic variation between domestic populations of plants and animals (Sánchez-Villagra et al., 2016) as well as a strong differentiation between wild and domestic populations. Both natural and artificial selection in domestic species have left footprints across the genome, known as selection signatures, which can point to regions harboring functionally important sequences of DNA (Qanbari and Simianer, 2014;Bertolini et al., 2018). Selection leads to changes in allele frequencies and haplotype patterns within populations under selection (Ma et al., 2015). Theoretically, a beneficial variant that has been under selection can show long-range linkage disequilibrium (LD) and high frequency over a long period of time (Grossman et al., 2010). Therefore, patterns of LD and variation in allele frequencies allow us to detect such selection signatures in genomes. When successful, selection signature identification can provide novel insights into mechanisms that create diversity across populations and contribute to mapping of genomic regions underlying selected traits or phenotypes (Andersson and Georges, 2004;Oleksyk et al., 2010). Approaches for detecting selection signatures rely on scanning variants across the genome of interest, searching for regions in which (1) the allele frequency spectrum is shifted towards extreme (high or low) frequencies; (ii) there is an excess of homozygous genotypes; (iii) there are long haplotypes with high frequencies and (iv) there is an extreme differentiation among populations. Several statistical methods has been developed to search for selection signatures, such as extended haplotype homozygosity (EHH) (Sabeti et al., 2002), integrated haplotype score (iHS) (Voight et al., 2006), hapFLK (Fariello et al., 2013), Cross Population Extended Haplotype Homozogysity (XP-EHH) (Sabeti et al., 2007), the composite likelihood ratio (CLR); runs of homozygosity (ROH) (Ceballos et al., 2018) and F_ST_ statistics (Weir et al., 2005).

In fishes, domestication is recent in comparison with other land animals (Teletchea, 2018), although some evidence of fish farming dates back approximately 3,500 years ago (Balon, 2004). An exponential development of aquaculture has occurred since 1960, relying on the domestication of a handful aquatic species (Teletchea, 2018). Salmonid species in particular have seen very intensive production increases over the last four decades. Currently, the most important farmed salmonids species in the word are Atlantic salmon (*Salmo salar*), rainbow trout (*Oncorhynchus mykiss*) and coho salmon (*Oncorhynchus kisutch*). Coho salmon belong to the *Oncorhynchus* genus and are naturally distributed across Pacific North American and Asian watersheds (Groot and Margolis, 1991). About 90% of farmed coho salmon production is based in Chile (<u>FAO, 2016</u>). Farmed populations of coho salmon were established in Chile at the end the 1970’s, with the importation of ova from the Kitimat river (British Columbia) and the US state of Oregon (Neira, 2014). During the 1990’s, a variety of breeding programs have been implemented for this species in Chile (Neira, 2014), mainly focused on improving growth, disease resistance traits and flesh color (Neira, 2014). The implementation of such breeding programs have likely, shaped the genetic diversity and haplotype structure of these populations, and their present genomes may contain traceable signatures of selection.

Genomic scans for detecting selection signatures have been successfully applied to several domestic animals, including aquaculture species. For instance, selection signature scans have been carried out for Atlantic salmon (Mäkinen et al., 2014;Gutierrez et al., 2016;Liu et al., 2016;López et al., 2018;López et al., 2019;Naval-Sanchez et al., 2020), Nile tilapia (Xia et al., 2015;Cadiz et al., 2019); and channel catfish (Nam et al., 2019). To date, there are no studies focusing on the identification of selection signatures in coho salmon populations. To increase the knowledge of domestication and selection effects on the genome diversity in farmed populations of this species, we searched for selection signatures in two breeding populations of coho salmon using different statistical approaches based in haplotype structure and allele frequency spectrum. Tracing signatures of selection in this species could provide further insights on genomic regions responsible for important aquaculture features of domestic Pacific salmon.

## Methods

### Populations

In this study we used two domestic populations from a breeding program of coho salmon maintained in Chile, namely Pop-A and Pop-B: we used 89 individuals for A and 43 individuals for B. These breeding populations were established in 1997 and 1998, belonging to even and odd spawning year, respectively (Yáñez et al., 2014). According to previous analyses (Ben Koop, unpublished data), it is very likely that these two populations correspond to a mixture of the two original broodstocks (Kitimat river and Oregon). Both populations have been selected to improve growth rate for around eight generations by using Best Linear Unbiased Prediction (Yáñez et al., 2016;Dufflocq et al., 2017) with genetic gains ranging between 6 to 13% per generation (Lhorente et al., 2019).

### Genotyping

Genomic DNA was extracted from fin clips of each individual and genotyped with a 200K Thermo Fisher Scientific Axiom® myDesign Custom Array developed by the EPIC4 genome consortium (Barria et al., 2019) (http://www.sfu.ca/epic4/).

Genotype calling was performed using Axiom Analysis Suite v3.1 (Thermo Fisher Scientific) following the Axiom Analysis user guide. After filtering, including removing markers with no position on the coho salmon reference genome (GCF_002021735.1) and markers that were identified as problematic (OTV, Call Rate Below Threshold, Other), 167,486 SNP were kept (Barria et al., 2019). In this study, we removed those markers that were not placed in chromosomes; therefore 136,500 markers were kept for downstream analysis. All these markers and samples passed the quality criteria (missing call rate ≤ 0.1).

### Genetic diversity, linkage disequilibrium (LD) and population structure

A common subset of 72,616 SNPs with MAF ≥ 0.05 and in Hardy-Weinberg equilibrium in both populations A and B were explored to in order to describe the genetic diversity of each population. Three parameters, including the number of SNPs with MAF ≥ 0.05; MAF ≥ 0.200 (N_SNP_), observed heterozygosity (H_O_), and expected heterozygosity (H_E_) were calculated using PLINK (Purcell et al., 2007).

We used the squared correlation coefficient between SNP pairs (*r^2^*) to measure LD (Hill and Robertson, 1968), which was calculated for all syntenic marker pairs. To enable a clear presentation of results showing LD in relation to physical distance between markers, SNP pairs were put into bins of 100 Kb, and mean values of *r*^*2*^ were calculated for each bin. The mean *r*^*2*^ for each of the distance bins was then plotted against the distance bin range (Mb). This analysis was carried out on a chromosome-by-chromosome basis.

To visualize genetic differentiation between populations a Principal component analysis (PCA) was conducted using PLINK. In addition, ADMIXTURE (Alexander et al., 2009) was employed to analyze the population structure, which was run with 10,000 iterations using the correlated allele model with K value from 1 to 20 to choose the optimal number of clusters at the lowest cross-validation error. Results were plotted using R (R core team, 2019).

### Selection signatures

We combined three methods (XPEHH, |iHS| and CLR) which have shown to have a power >70% to detect selection signatures in comparison with other combinations of statistical tests (Ma et al., 2015).

1. **XP-EHH.** While, the iHS statistic compares the integrated EHH profiles between two alleles at a given SNP in the same population (Voight et al., 2006), the XP-EHH (Cross Population Extended Haplotype Homozygosity) statistic compares the integrated EHH profiles between two populations at the same SNP (Sabeti 2007). Therefore, the computation of EHH is required for each population. The XP-EHH statistic is then defined as ln(I_PopA_/I_PopB_), where I_PopA_ is the integrated EHH for the population A and I_PopB_ is the integrated EHH for population B. Negative XP-EHH scores suggest selection in B, whereas positive scores suggest selection acting in the A. A −log_10_(*p* value) of three (*p* value ≤ 0.001) was used as the threshold for considering XP-EHH score as significant evidence of selection and at least two SNPs ≤ 500 kb apart.
2. **iHS.** The iHS statistics was used to detect footprints of selection within the studied populations. This test is based on the standardized log ratio of integrals of the decay of the extended haplotype homozygosity (EHH), computed for both ancestral and derived alleles of the focal SNP. Selection induces hitchhiking (genetic draft), that leads to extended haplotypes for the selected allele and a slower fall-off of EHH on either side of the selected locus. Thus, a high iHS scoring SNP is typically associated with longer haplotypes and lower neighborhood diversity compared to the other SNPs (Wagh et al., 2012). Phased haplotypes were obtained using Beagle (Browning and Browning, 2009). The ancestral and derived alleles of each SNP were inferred in two ways: (i) the SNP probes were compared with the genome of Atlantic salmon, which was used as an outgroup species. (ii) for SNPs that could not be compared, the ancestral allele were inferred as the most common allele in the total dataset, as suggested in other studies (e.g. Bahbahani et al., 2015; Hacia et al., 1999). The iHS score was computed for each autosomal SNP using the R package rehh 3.1.1 (Gautier et al., 2017), using default options.
3. **CLR**: We implemented CLR test in the software SweeD V3.3.2 (Pavlidis et al., 2013). SweeD evaluates the variation in the site-frequency spectrum along the chromosome and implements the composite likelihood ratio (CLR) statistics (Nielsen et al., 2005). CLR computes the ratio of the likelihood of a selective sweep at a given position to the likelihood of a null model without a selective sweep. We calculated the CLR within each population and for each chromosome separately at grid points for every 20 Kb. We used the 99.5th percentile of the distribution of CLR scores as threshold for the detection of outliers.

### Candidate genes and functional analysis

A genomic region was considered as being under selection if it matched the criteria described above for iHS, XP-EHH and CLR. For each identified selective sweep region, we extended the region containing outlier scores by adding 250Kb to each end, to account for potential blocks of high linkage disequilibrium. Gene annotation was performed using GCF_002021735.1 coho salmon genome assembly from the NCBI Eukaryotic genome annotation pipeline. .

The Database for Annotation, Visualization, and Integrated Discovery (DAVID) v6.8 tool (Huang et al., 2009) was used to identify Gene Ontology (GO) terms and KEGG (Kyoto Encyclopedia of Genes and Genomes) pathways using a list of genes within significant regions based on iHS, XP-EHH, CLR values and the zebrafish annotation file as a reference genome.

## Results

Population B has a slightly greater H_O_ = 0.39 than Pop-A H_O_ = 0.37. The number of SNPs with MAF ≥ 0.05 was higher in Pop-A than Pop-B. In addition LD presented a higher level in Pop-B (r^2^ = 0.14) than Pop-A (r^2^ = 0.10) (Table 1). LD decayed faster in Pop-A, reaching a level of 0.14 around 500 Kb, while in Pop-B at the same distance LD reaches a value of 0.18. A similar pattern was observed for most of the chromosomes, except for, and Oki-01, Oki-21, Oki-28, where LD decay was slightly higher in Pop-B and Oki-10, where both populations presented a similar pattern of LD decay (Supplementary Fig. S1).

**Table 1.**
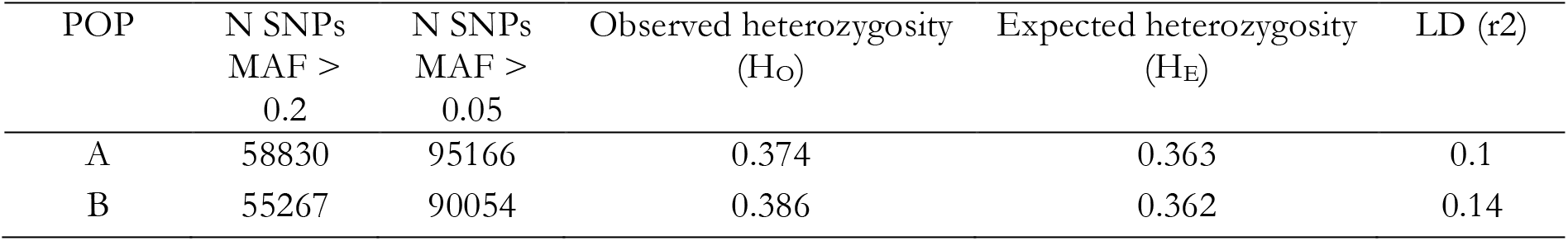
Basic statistics in term of number of SNPs greater to MAF> 0.05 or 0.2; Observed heterozygosity (H_o_); Expected heterozygosity (H_E_) and LD level across two coho salmon populations.

Principal component analysis (PCA) was employed to explore the clustering of individuals from the two populations. The PCA revealed two distinct clusters corresponding to population A and B. Principal components (PCs) 1, 2 and 3 jointly accounted for 39.7% of total variance, with PC 1 (26.1%) separating A and B; PC 2 (7.6%) and PC 3 (6%) not clearly discriminating populations. However, Pop-B produced a much more heterogeneous group with individuals spread along both PC 2 and 3 (Fig. 1). ADMIXTURE analysis revealed that the lowest cross-validation error was reached at K=10 (Supplementary Fig. S2), showing 10 ancestral lineages for these two populations (Fig. 2).

**Figure 1.**
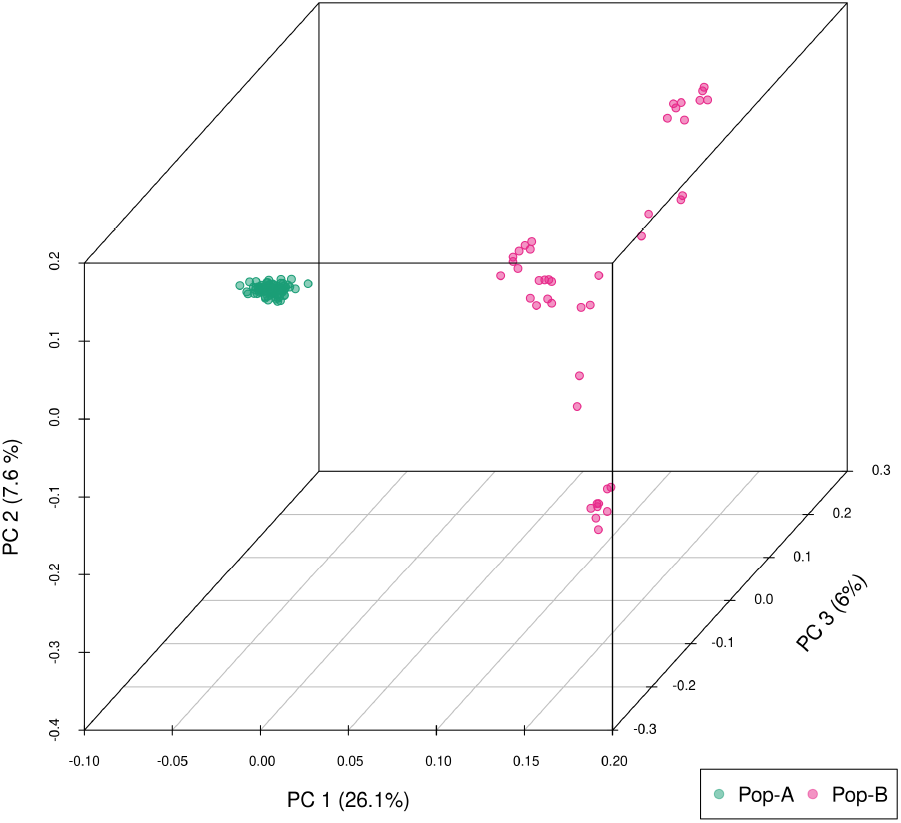
Principal components analysis (PCA) of genetic differentiation among individuals. Three first nents are shown. Each point represents one individual, and different colors represent populations: Pop-n, Pop-B = magenta.

**Figure 2.**
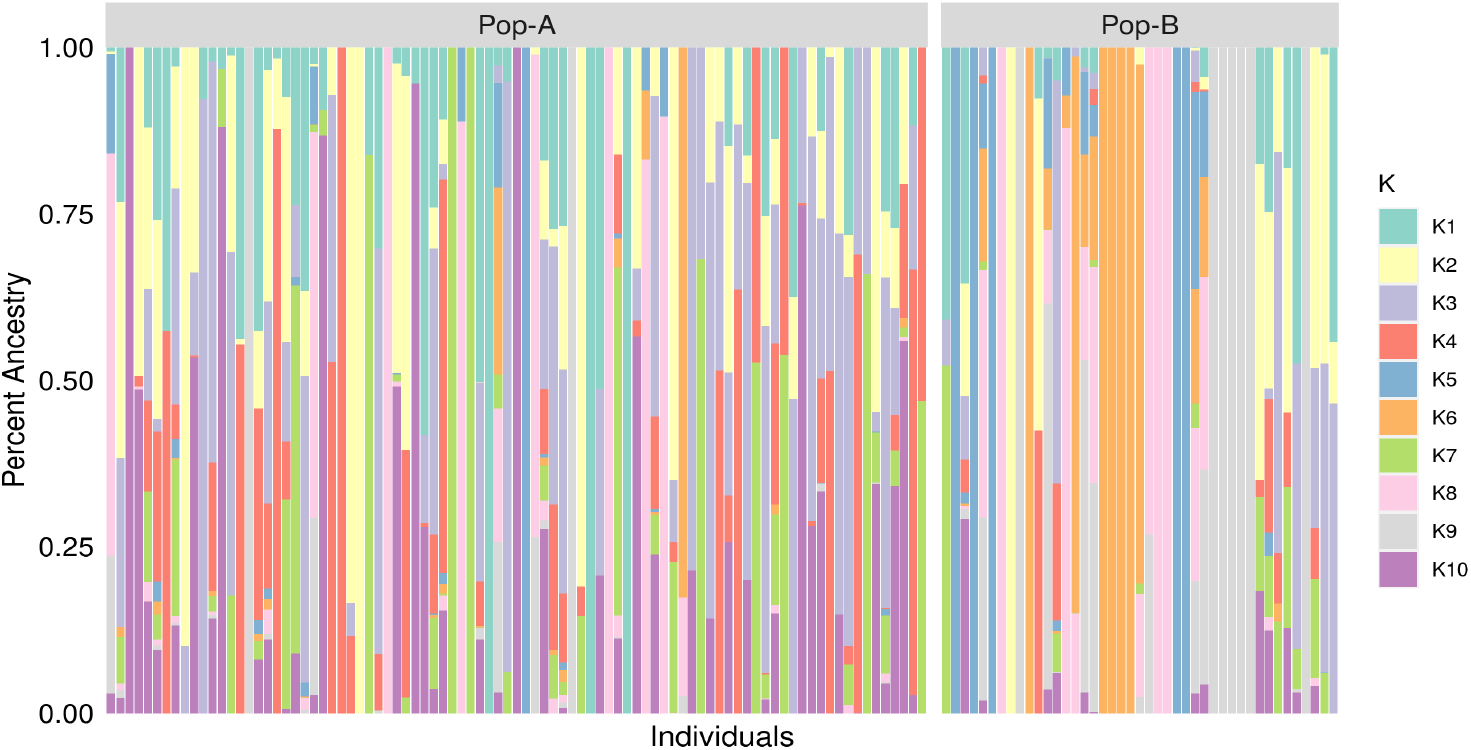
Individual assignment probabilities generated with ADMIXTURE (1⩽K⩽20). Each color represents a and the ratio of vertical lines is proportional to assignment probability of and individual to each cluster.

**Figure 3.**
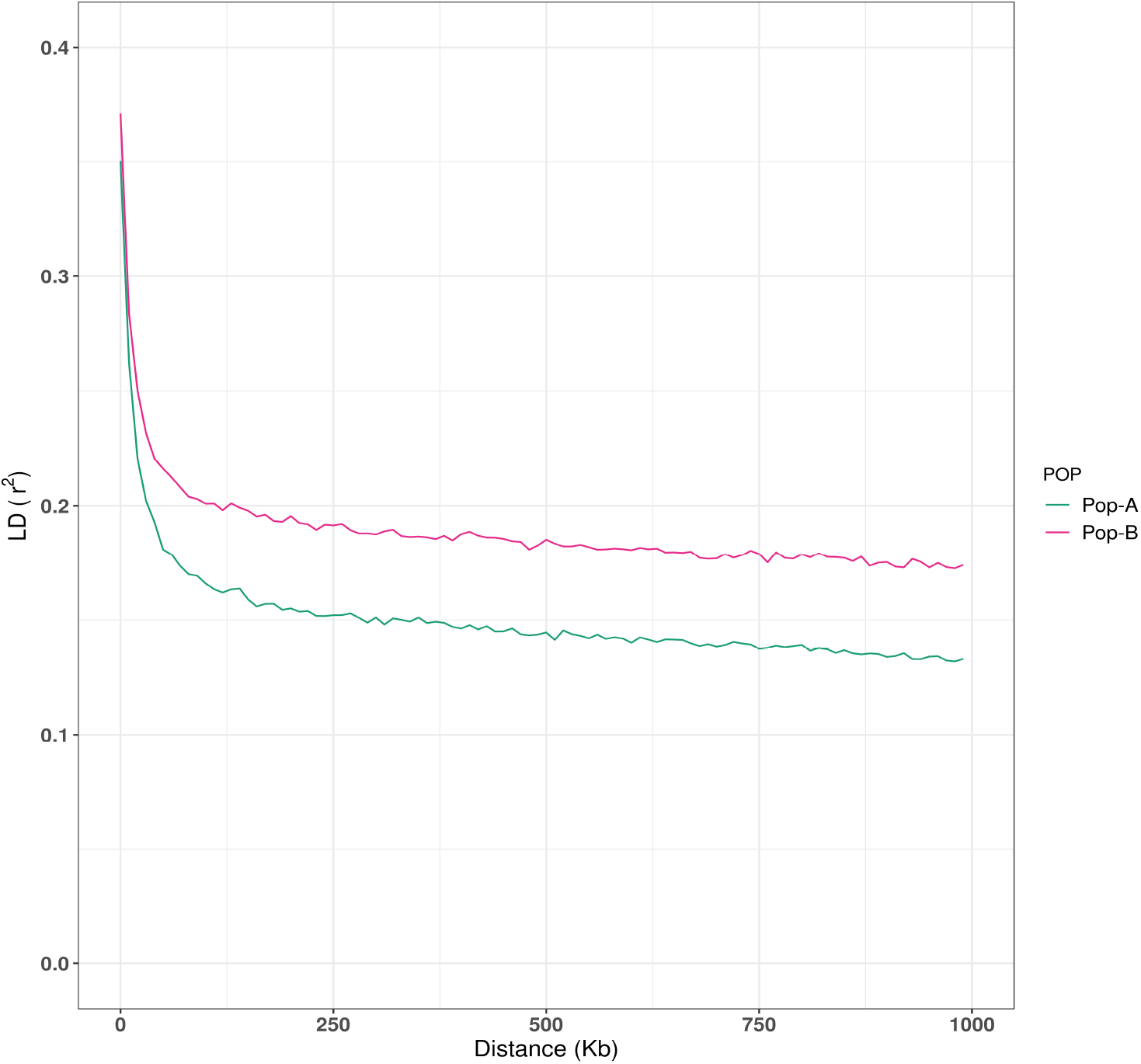
Decay of average linkage disequilibrium (r^2^) over distance across the two farmed populations. Different nes represent populations: Pop-A=green, Pop-B = magenta.

### Genome-wide scanning for selection signatures

To detect signatures of selection in the two coho salmon breeding populations, we used three statistical methods. Two of them are based in extended haplotype homozygosity (EHH): iHS and XPEHH, and one method is based on the variation of the site-frequency spectrum, the composite likelihood ratio (CLR) tests.

Inner circles of Figures 4 and 5 show the genome-wide distribution of **iHS** values. The iHS statistics revealed 156 and 99 SNPs with significant iHS scores (*p* ≤ 0.001) for populations A and B respectively. We found 26 and 16 regions in these respective populations with at least two significant SNPs that are ≤ 500 Kb apart. These genomic regions spanned 17.1 Mb across chromosomes Oki5, Oki6, Oki28 and Oki29 in Pop-A, and 7.5 Mb in Pop-B in Oki5, Oki14, Oki15, Oki17, Oki18, Oki22 and Oki26 (Table S1). The highest values were present in Oki5 and Oki28 for Pop-A, and Oki14 and Oki18 for Pop-B. **XP-EHH** revealed 43 SNPs, which surpassed the significance threshold (Figures 4.B and 5.B), 41 with positive scores and indicating selection in Pop-A, and two with negative scores indicating selection in Pop-B (Table S2). According to the criteria that we defined, six genomic regions spanning 3.72 Mb were detected in Pop-A, and one region of 520.8 Kb in Pop-B. Genomic regions with XP-EHH values above 3 were detected in Oki1, Oki10 and Oki28 for Pop-A. The highest XP-EHH estimate was on Oki1 (XP-EHH= 3.8030, −log_10_(p-value)= 3.8448), and the lowest value was on Oki15 (XP-EHH=−3.373854, log_10_(p-value)=3.130043). As a complementary method to iHS and XP-EHH, we performed the composite likelihood ratio (**CLR**). Figures 4.C and 5.C illustrate the CLR statistics against the genome positions for coho salmon. Based on a conservative 99.9% outlier threshold (CLR_A_ ≥ 4.81, CLR_B_ ≥ 4.64), we detected 95 regions affected by selective sweeps, covering 28.24 Mb and 21.92 Mb in population A and B, respectively (Table S3). In both populations, the results provided evidence of selective sweeps in several chromosomes across the genome. In Pop-A the highest CLR values were located on Oki14 (CLR = 9.98) within a region of 180 Kb and on Oki17 (CLR = 9.35) within a region of 400Kb. While, in Pop-B a clear evidence of selective sweep is observed on Oki15 (CLR = 10.75), with a cluster of extreme signals in a genomic region of 780Kb.

**Figure 4.**
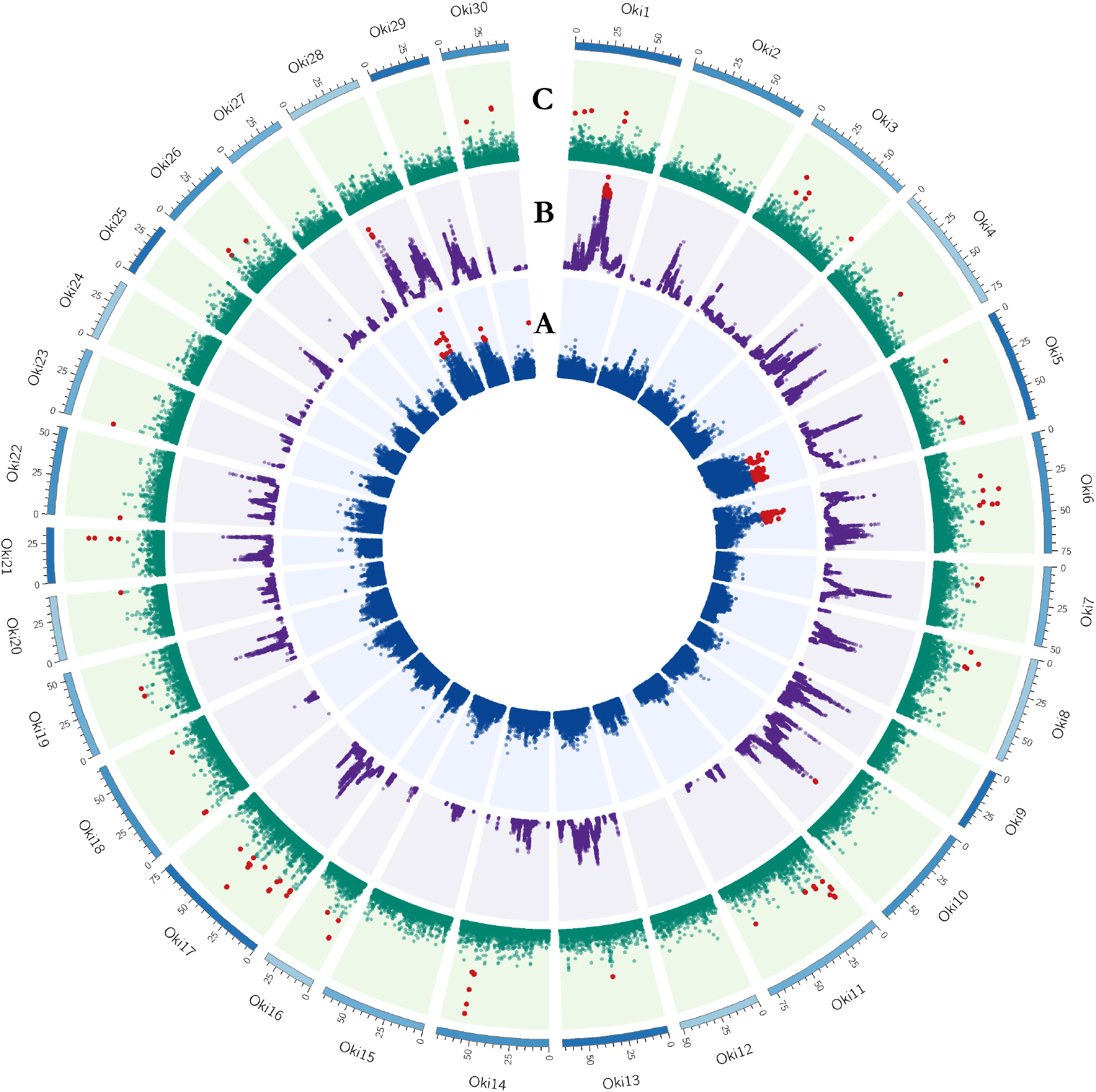
Circos plot of the global distribution of selection signatures across the genome in Pop-A. The circles from inside to outside illustrate: **A.** −log_10_(p-value) of iHS (Blue); **B.** −log_10_(p-value) of XP-EHH (Purple) and **C.** CLR values (Green). Values for each test surpassing the threshold (iHS and XP-EHH ≥ 3, and CLR ≥ 4.81) are highlighted in red.

**Figure 5.**
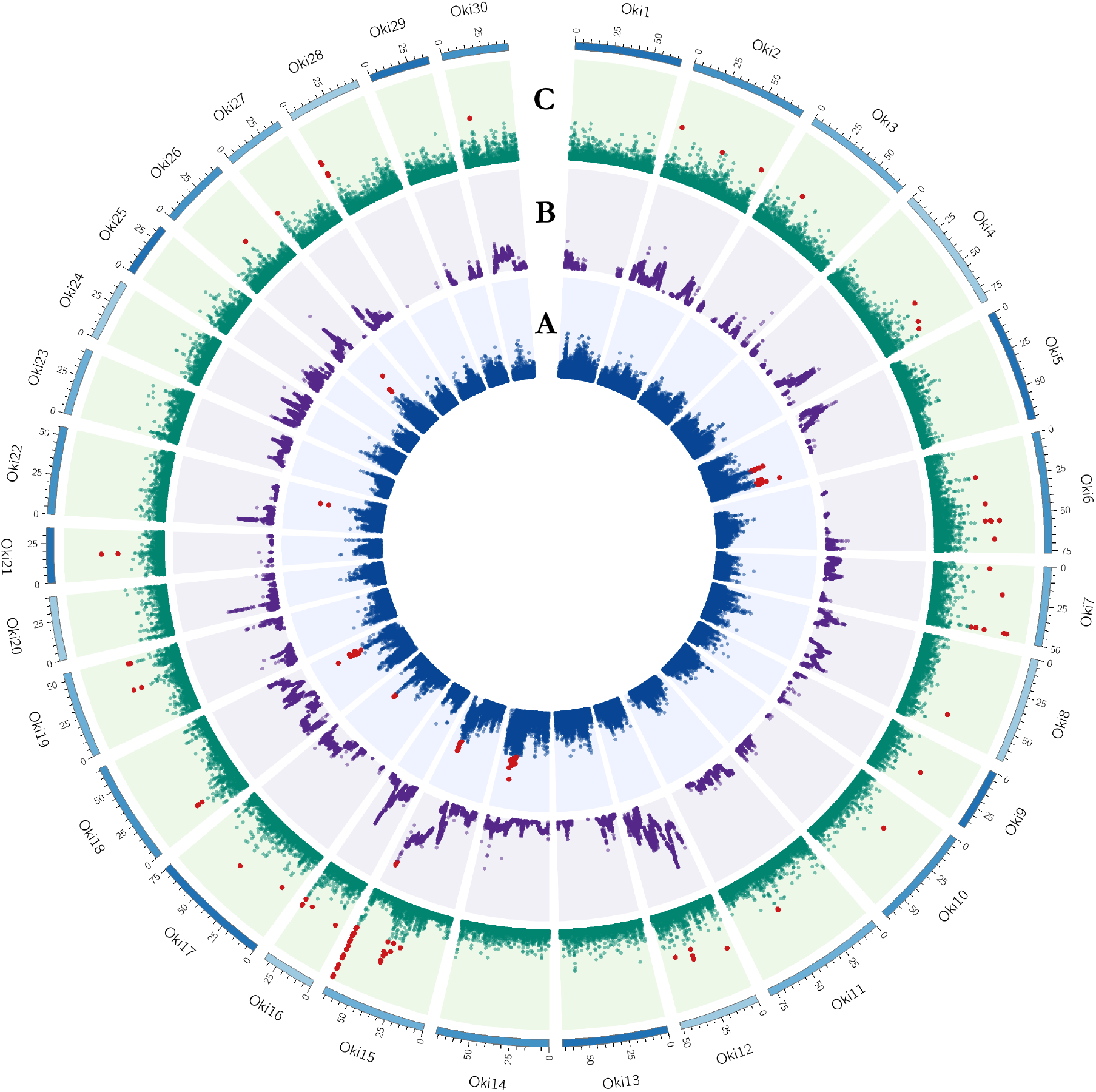
Circos plot of the global distribution of selection signatures across the genome in Pop-B. The circles from inside to outside illustrate: **A.** −log_10_(p-value) of iHS (Blue); **B.** −log_10_(p-value) of XP-EHH (Purple) and **C.** CLR values (Green). Values for each test surpassing the threshold (iHS and XP-EHH ≥ 3, and CLR ≥ 4.64) are highlighted in red.

### Functional characterization of genomic regions

A total of 1118 and 812 genes within the identified genomic regions putatively subjected to selection were retrieved for Pop-A and Pop-B, respectively. Gene annotation of selected regions revealed several functionally important genes, such as *kdm6a* involved in the regulation of anterior/posterior body axis development in zebrafish (Lan et al., 2007) or *sec24d* and *robo1* which were associated to resistance against *Piscirickettsia salmonis* in farmed coho salmon populations from Chile (Barría et al., 2018). Genes in genomic regions with the highest scores for iHS, XP-EHH and CLR are shown in Tables 2–4 while the complete list of the genes found is presented in Table S4.

**Table 2.**
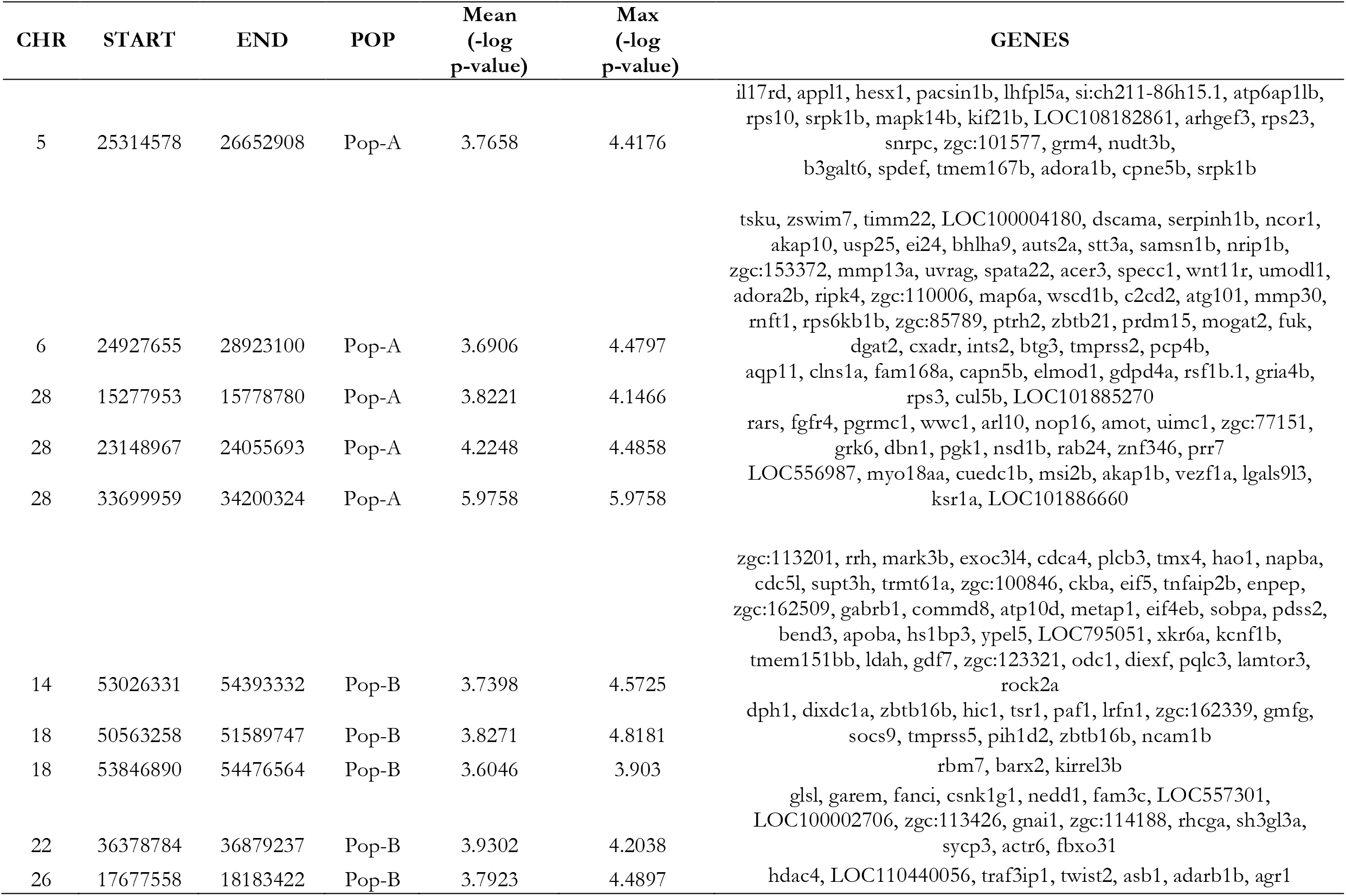
Genomic regions and genes spanning the five strongest detected selection signatures by iHS in each population.

**Table 3.**
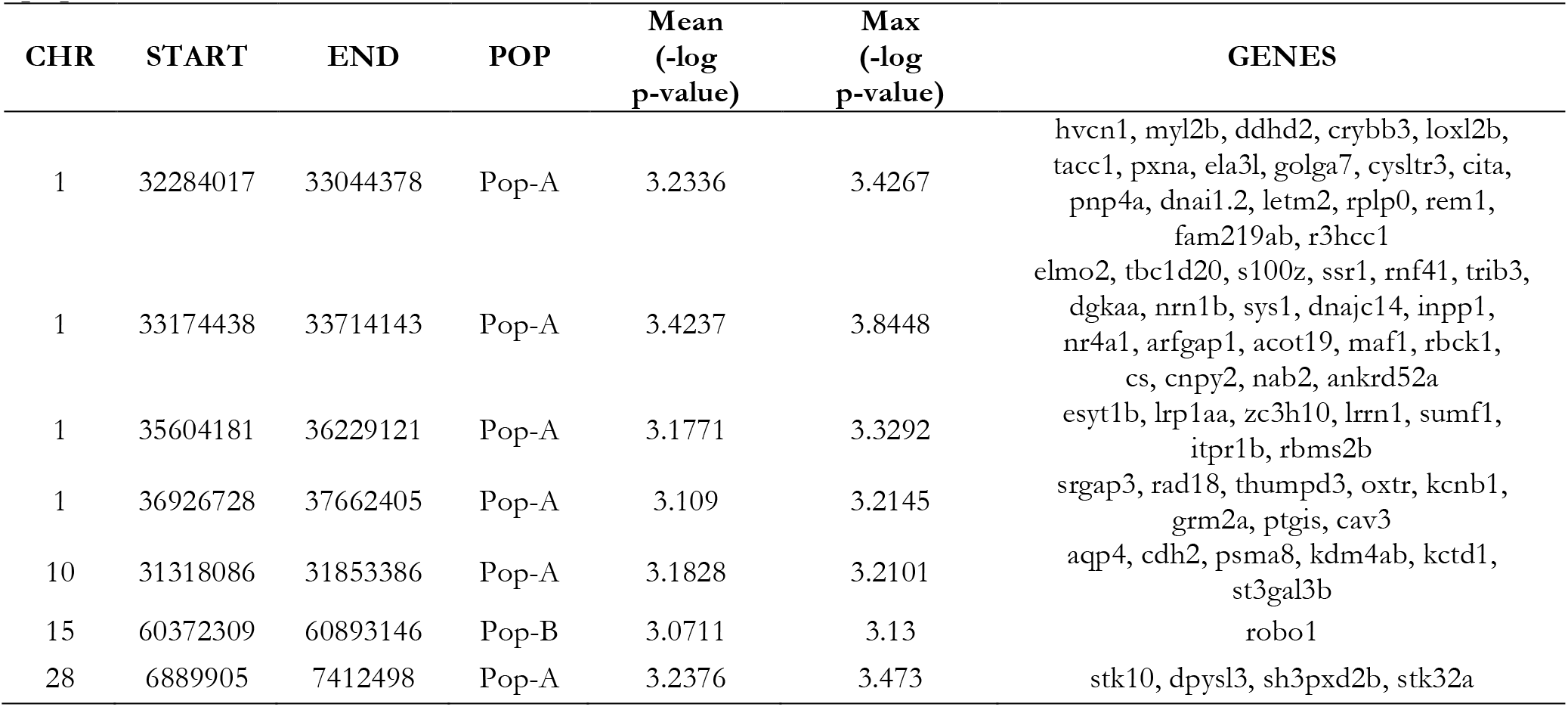
Genome regions and genes spanning the detected selection signatures by XP-EHH in each population.

**Table 4.**
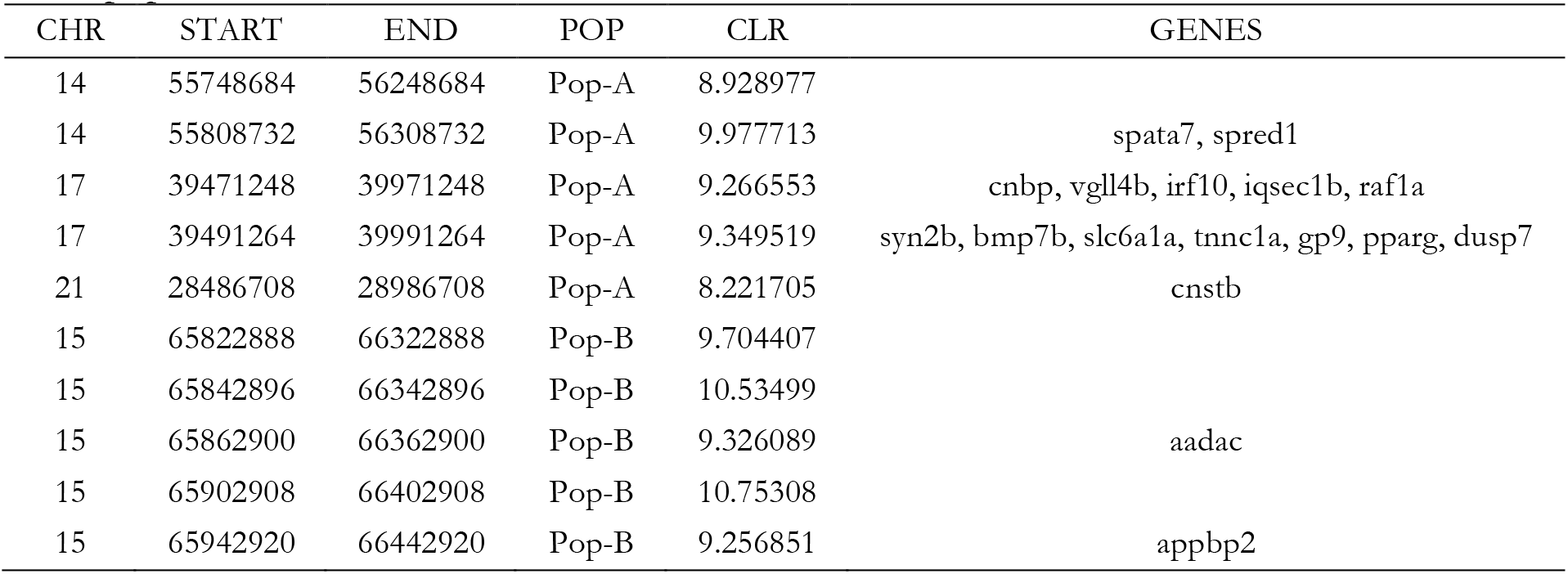
Genome regions and genes spanning the five detected selection signatures by CLR in each population.

We further investigated the functions associated with the putative genes undergoing positive selection by analyzing over-represented GO terms and pathways using DAVID. The identified GO terms and pathways are shown in Table S5. A total of 21 and 13 GO terms were observed for A and B, respectively. This includes 13 terms for biological process (BP), 3 for molecular function (MF), and 9 for cellular component (CC). The BP category is shown in Figure 6; most of these terms are associated with basic metabolic processes, such as Developmental Process, Multicellular Organismal Process, Single-Organism Process, Response To Stimulus, Developmental Process, Presynaptic Process and Growth, among other. Additionally, 11 pathways were enriched (Table S5) such as Vascular smooth muscle contraction, Endocytosis, Regulation of actin cytoskeleton and TGF-beta signaling pathway

**Figure 6.**
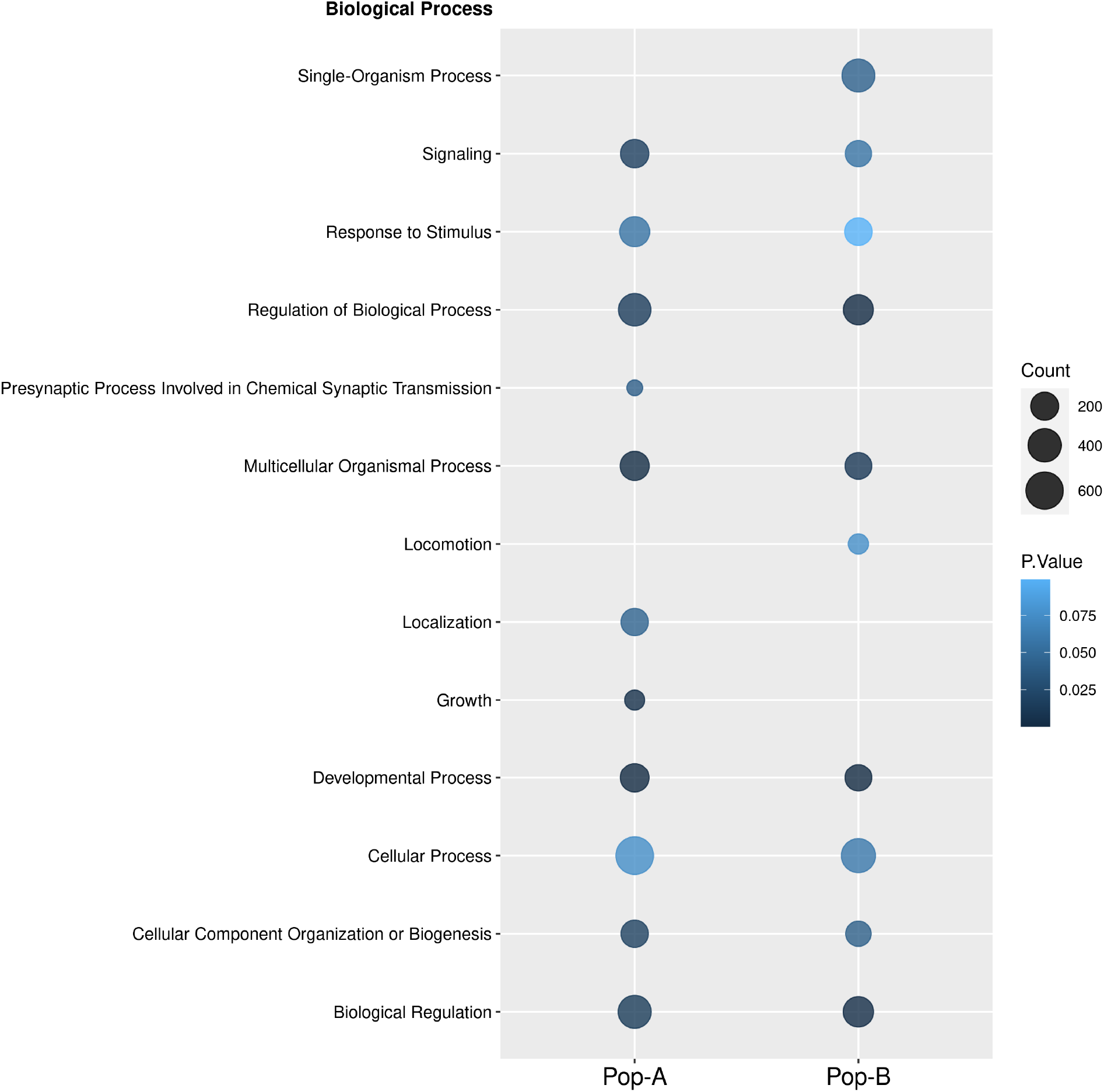
Functional annotation of candidate selective sweeps genes. Gene ontology (GO) analysis performed for al process. Color intensity of circle shows the significance of GO term and the size of the circle represents ber of genes associated with the GO term.

## Discussion

Selective events are the major mechanism driving differentiation of populations. In this study, we assessed the genetic structure and detected selection signatures in two domestic populations of coho salmon by using three complementary statistical methods. First, we investigated basic population genetics statistics. Genetic diversity estimates, assessed by observed and expected heterozygosity, were in the upper limit of those reported in other salmon species introduced for farming purposes in Chile, such as Atlantic salmon, in which heterozygosity levels of these populations have been shown to range from 0.266 to 0.37 (López et al., 2018;López et al., 2019). In addition LD, which can be informative of population demography, decays similarly to what has been observed on previous studies in farmed coho salmon population from Chile (Barria et al., 2019), and more rapidly than farmed Atlantic salmon (Barria et al., 2018). Thus, the higher levels of genetic diversity and the lower levels of LD in farmed coho salmon populations suggest less impact of artificial selection on genome diversity, likely due to a shorter period of time of domestication in comparison with Atlantic salmon aquaculture populations from Chile.

Several methods have been developed for detecting selection signatures in the genome. These methods can be grouped into three categories based on the type of genetic information exploited: population differentiation, site-frequency spectrum and linkage disequilibrium approaches (Oleksyk et al., 2010). In the present study we implemented three different tests (iHS, XP-EHH and CLR) to comprehensively identify candidate regions of positive selection in coho salmon. As is shown in Figure 8, different genomic regions were detected by these methods. CLR have the largest overall number of signals detected to be underlying selection, followed by iHS and XP-EHH. No overlap was observed among the three methods, however CLR and iHS (Fig. 7), showed five overlapping regions on Oki5, Oki6 and Oki14 spanning a total of 2.02 Mb. Low overlap among methods based on allele frequencies compared to those based on haplotype patterns has been observed in previous studies in Atlantic salmon (Mäkinen et al., 2014; López et al., 2018). Discrepancies between loci under putative selection detected by different methods are expected, given that they benefit from different sources of information from genome variation. For instance, iHS test has advantage in detecting selective sweep with variants at intermediate frequencies (Sabeti et al., 2007), while XP-EHH is more powerful at detecting complete or nearly complete selective sweeps (Sabeti et al., 2007).

**Figure 7.**
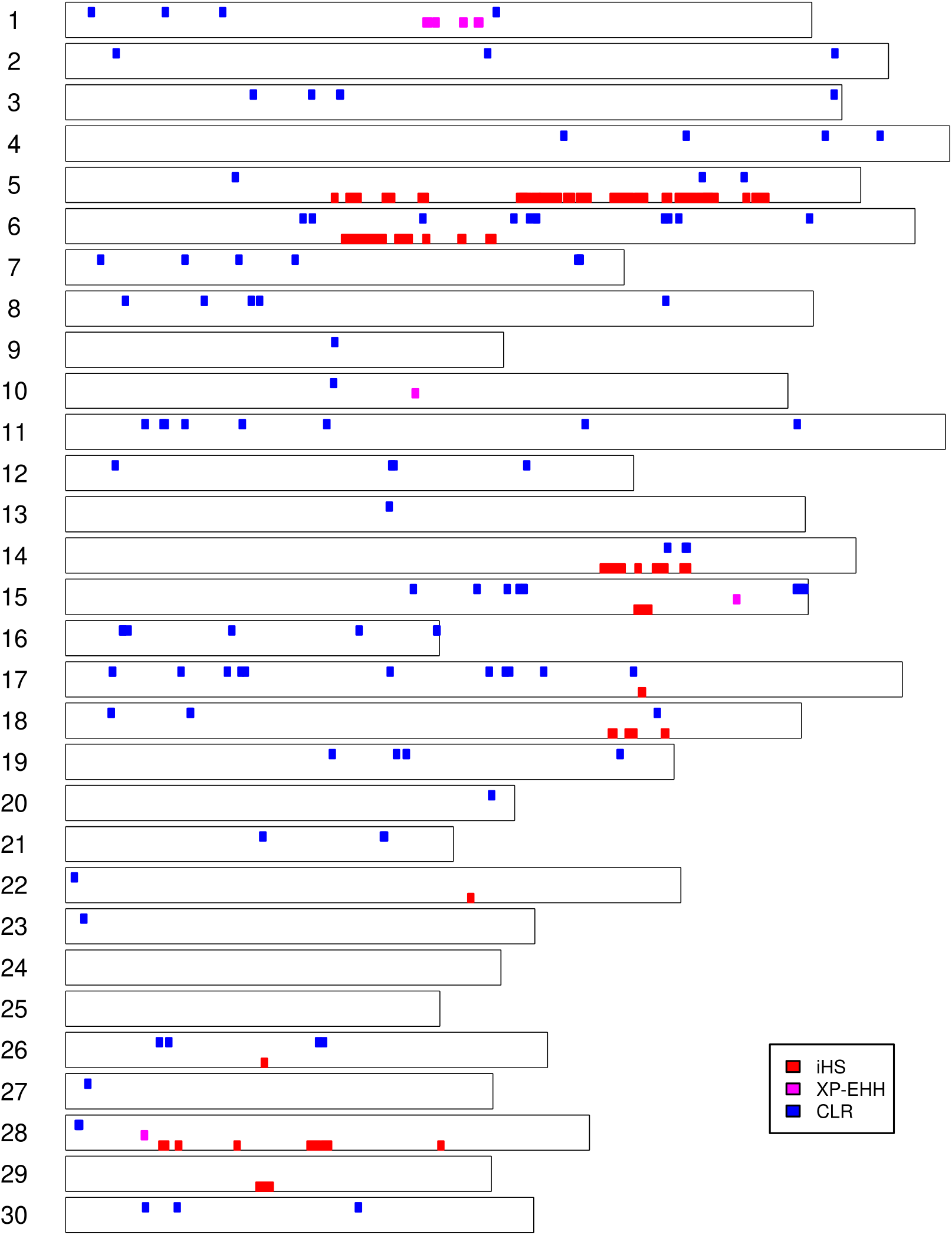
Genomic distribution of selection signatures detected by iHS, XP-EHH and CLR on all chromosomes.

**Figure 8.**
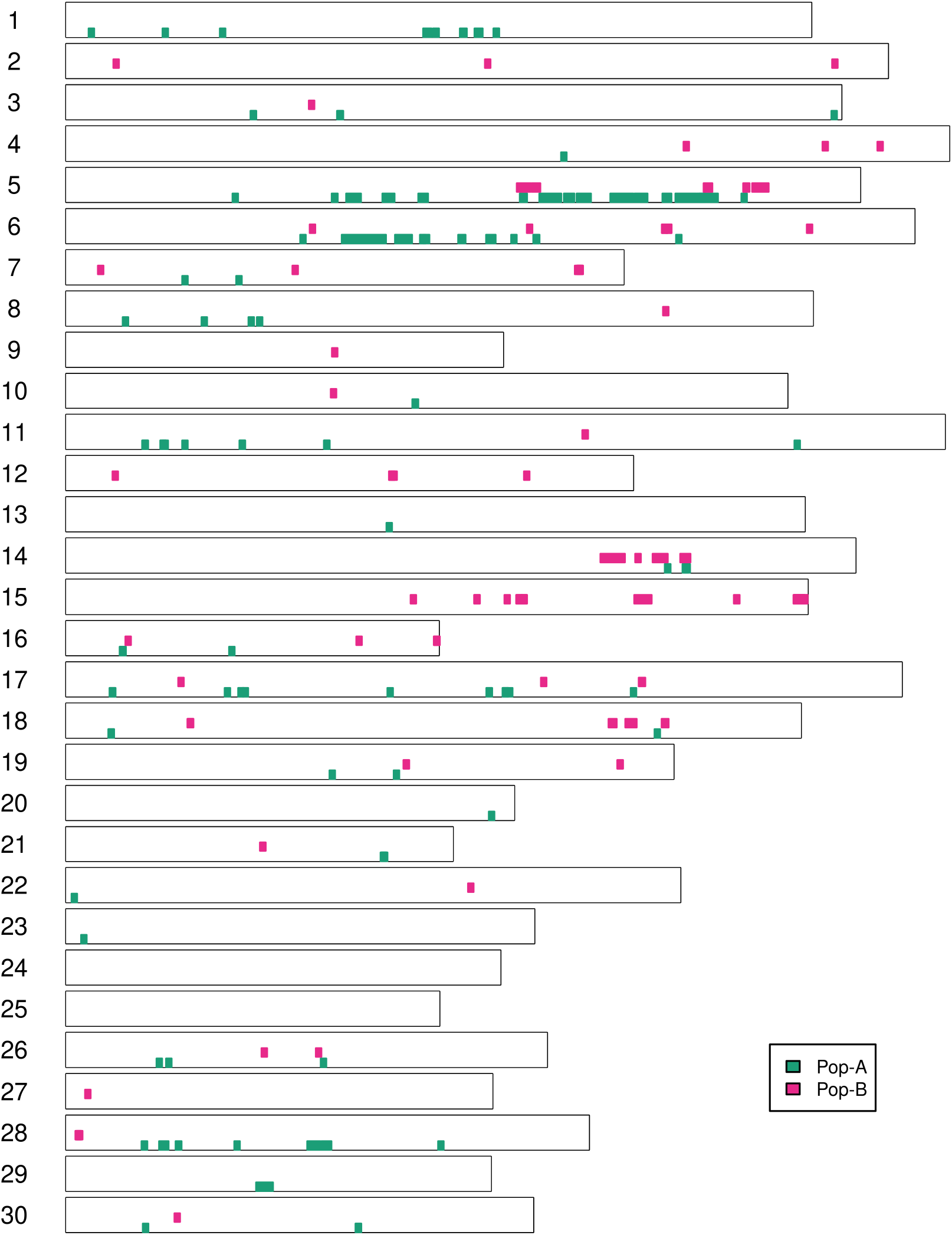
Genomic distribution of selection signatures detected per populations on all chromosomes.

On the other hand, a low overlap between the two populations was observed (Fig. 8). Four chromosomes showed common regions harboring selection signatures: Oki5 (1.86 Mb); Oki14 (946 Kb), Oki16 (41.5 Kb) and Oki26 (57.3 Kb). We suggest this low overlap among populations might be the effect of breeding program goals, which predominantly are focused on improving polygenic traits (e.g., growth), therefore, different genomics regions on different populations are affected by selection across the genome.

One of the goals of this study was to identify putative candidates genes involved in the domestication process and artificial selection of coho salmon. In accordance with the primary genetic improvement goal, where growth rate has been the main focus, we found a series of genes potentially relevant for this trait, including *anapc2*; *alad*; *bdkrb2*; *fam98b*; *chp2*; *myn* that were previously associated with body weight in a genome-wide association study in Atlantic salmon (GWAS) (Yoshida et al., 2017) and *sytl5, txnrd1* genes associated with growth traits in juvenile farmed Atlantic salmon (Tsai et al., 2015). The *chp2* gene was also found to be putatively under selection in an Atlantic salmon domestic population selected for fast growth (López et al., 2019). We also identified the *kdm6a* gene, which has been involved in the regulation of anterior/posterior body axis development in zebrafish (Lan et al., 2007) and has been related to growth-related traits in pigs (Guo et al., 2019). Furthermore, we identified the biological processes “Developmental Process” and “Growth”, within the significant terms associated with the genes harbored in the identified genomic regions, suggesting that loci controlling growth and development are most likely under selection in these farmed coho salmon populations.

On the other hand, host-pathogen interactions lead to strong selection in the genome of vertebrate species (Early and Clark, 2017). We suggest that adaptation to farming environment has imposed selection on genomic regions related to the immune system in coho salmon. Here we found the *cnpy2, synrg* and *med10* genes, which were previously shown to be affected by parasite-driven selection in Atlantic salmon (Zueva et al., 2014). More importantly, we detected the genes *sec24d* and *robo1* that have been associated with disease resistance in the face of a bacterial disease (*Piscirickettsia salmonis*) in coho salmon populations farmed in Chile (Barría et al., 2018).

Artificial selection may be applied either inadvertently (unconsciously) or intentionally (consciously) (Price, 2002). For most livestock species, it is thought that the early stages of domestication involved unconscious selection for behavioral traits (e.g., for tameness and reduced aggression), followed by selection focused on breeding objectives (Gregory, 2009). In fish, behavioral traits such as swimming capacities, foraging, social interactions, reproduction, or personality and cognitive abilities, are also modified by domestication (Pasquet, 2018). In the present study we identified the genes *robo1* and *dcdc2* which are associated to complex cognitive acquired skills, including spoken and written language in human (Mozzi et al., 2016). We also identified the *gtf2ird1* associated to aggression and social interaction in mice (Young et al., 2008) and recently shown to be under selection in Nile tilapia (Cadiz et al 2020). We suggest that such genes could be associated with behavioral traits in coho salmon as well.

Sexual maturation and reproductive traits have profound importance from both evolutionary and economic perspective. In fish farming, maturation is frequently delayed by exposing fish to continuous light or light regimes, which are different to those, found in natural environment, affecting the perception of seasonality and circannual rhythms (Taranger et al., 2010). In this study, we identified the *picalmb* and *tsku* genes with evidence of selection. These genes were recently associated to maturation traits in Atlantic salmon (Mohamed et al., 2019). Furthermore, we also detected the *vgll4b,* which has previously shown signs of selection in Atlantic salmon farmed populations (López et al., 2019). It is important to mention that a gene from the VGLL (*vgll3*) family is associated with age to maturity in natural populations of Atlantic salmon (Ayllon et al., 2015;Barson et al., 2015).

## Conclusion

This study reported the identification of selection signatures in the genome of aquaculture lines of coho salmon. Our analyses of two domestic populations revealed putative selection signatures of genomic regions containing genes involved in growth, immune system, behavior and maturation traits. As expected, these findings are congruent with the breeding goal for the populations studied here, which includes the most highly selected trait, growth rate. In addition, the effect of domestication and adaptation to captive environment, most likely affecting loci controlling traits that have not been directly part of the breeding objectives, including immunity and, behavior. These results provide further insights into the genome diversity changes driven by domestication and selection mechanisms in farmed coho salmon populations.

## Availability of data and material

We will archive the genotypic data from this study at FigShare once the manuscript is accepted for publication.

## Acknowledgments

We thank the Government of Canada through Genome Canada, Genome British Columbia, and Genome Quebec for supporting this research that was carried out in conjunction with the project EPIC4 (Enhanced Production in Coho: Culture, Community, Catch). We also would like to acknowledge Pesquera Antares S.A., which provided the fish used in this study. JMY is supported by Núcleo Milenio INVASAL funded by Chile’s government program, Iniciativa Científica Milenio from Ministerio de Economía, Fomento y Turismo.

## Authors’ contributions

MEL and JMY conceived and designed the study. MEL performed the analyses and wrote the the manuscript. MIC contributed with data analyses. MIC, EBR, BFK and JMY contributed to discussion and writing. All authors have reviewed and approved the final manuscript.

## Ethics approval and consent to participate

Coho salmon individuals and sampling procedures were approved by the Comité de Bioética Animal from the Facultad de Ciencias Veterinarias y Pecuarias, Universidad de Chile (certificate N° No. 08-2015).

## Consent for publication

Not applicable

