## Supplementary_legends for "Detection of selection signatures in farmed coho salmon (*Oncorhynchus kisutch*) using dense genome-wide information"

**Supplementary information**

**Supplementary Table S1.** Regions with at least two SNPs above the threshold, detected by iHS. Definition of columns: **(1)** Chromosome **(2)** Position of the first SNP **(3)** Position of the last SNP **(4)** Start of the region (First SNP minus 250Kb) **(5)** End of the region (Last SNP plus 250 Kb) **(6)** Region name **(7)** Maximum –log_10_(p-value) in the region **(8)** Maximum |iHS| in the region **(9)** Number of SNPs in the region **(10)** Population.

**Supplementary Table S2.** Regions with at least two SNPs above the threshold, detected by XP-EHH. Definition of columns: **(1)** Chromosome **(2)** Position of the first SNP **(3)** Position of the last SNP **(4)** Start of the region (First SNP minus 250Kb) **(5)** End of the region (Last SNP plus 250 Kb) **(6)** Region name **(7)** Maximum –log_10_(p-value) in the region **(8)** Maximum XP-EHH in the region **(9)** Number of SNPs in the region **(10)** Population.

**Supplementary Table S3.** Regions above the threshold, detected by CLR. Definition of columns: **(1)** Chromosome **(2)** Start of the region (First SNP minus 250Kb) **(3)** End of the region (Last SNP plus 250 Kb) **(4)** Region name **(5)** ALPHA value in the region **(6)** CLR score in the region **(7)** Population.

**Supplementary Table S4.** Genes identified by iHS, XP-EHH and CLR. Definition of columns: **(1)** Chromosome **(2)** Start of the region **(3)** End of the region **(4)** Population **(5)** Test **(6)** Gene Name in Coho salmon **(7)** Gene Name in zebra fish.

**Supplementary Table S5.** Gene Ontology (GO) terms and KEGG (Kyoto Encyclopedia of Genes and Genomes) pathways identified in this study based on iHS, XP-EHH, CLR results.

**Supplementary Figure S1.** **.** Decay of linkage disequilibrium (LD) by chromosome for each population. Different color lines represent populations: Pop-A=green, Pop-B = magenta.

**Supplementary Figure S2.** Cross-validation error for ADMIXTURE results calculated for K values from 1 to 20.
