## Supplementary figures and images for "Detection of selection signatures in farmed coho salmon (*Oncorhynchus kisutch*) using dense genome-wide information"

### Figure S1

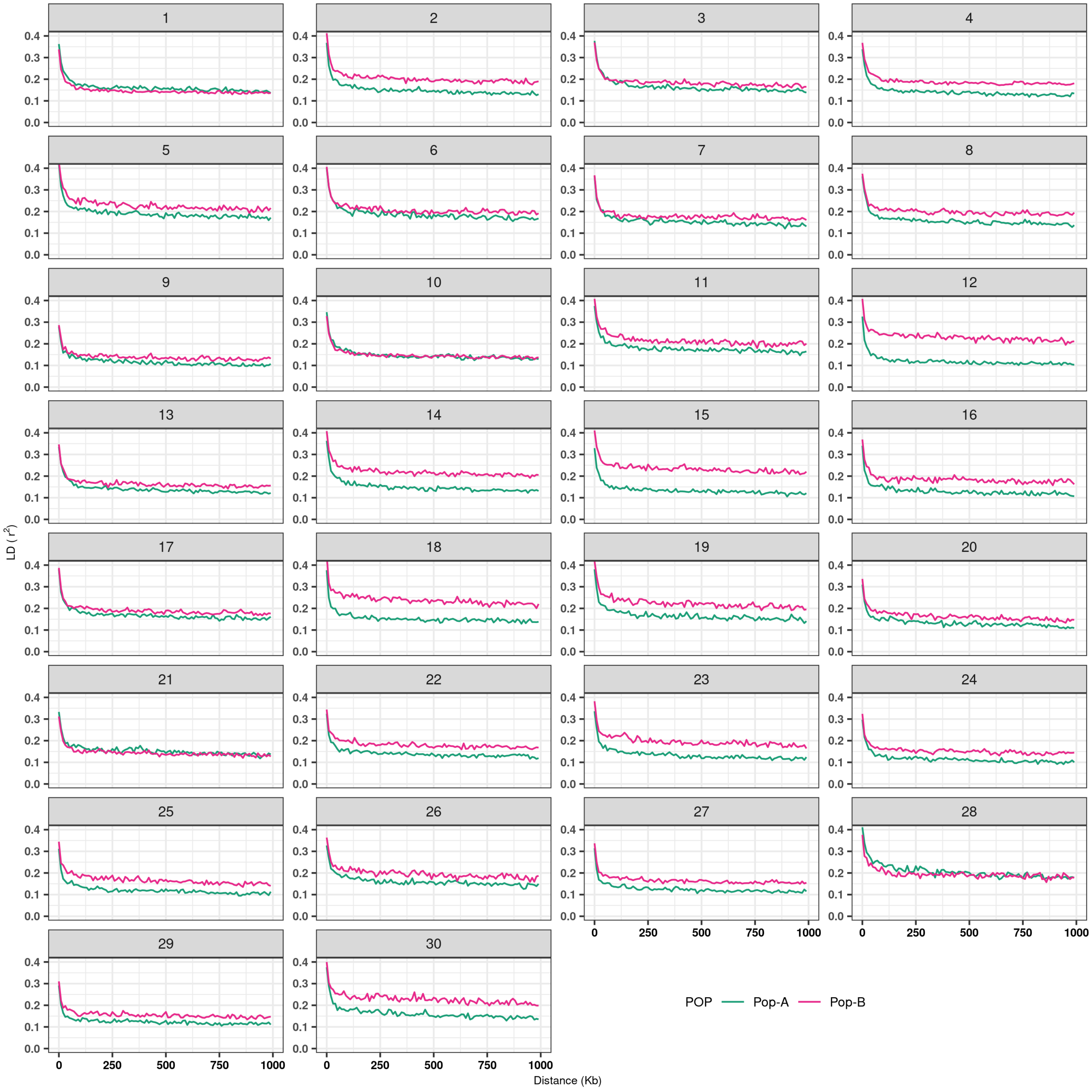

### Figure S2

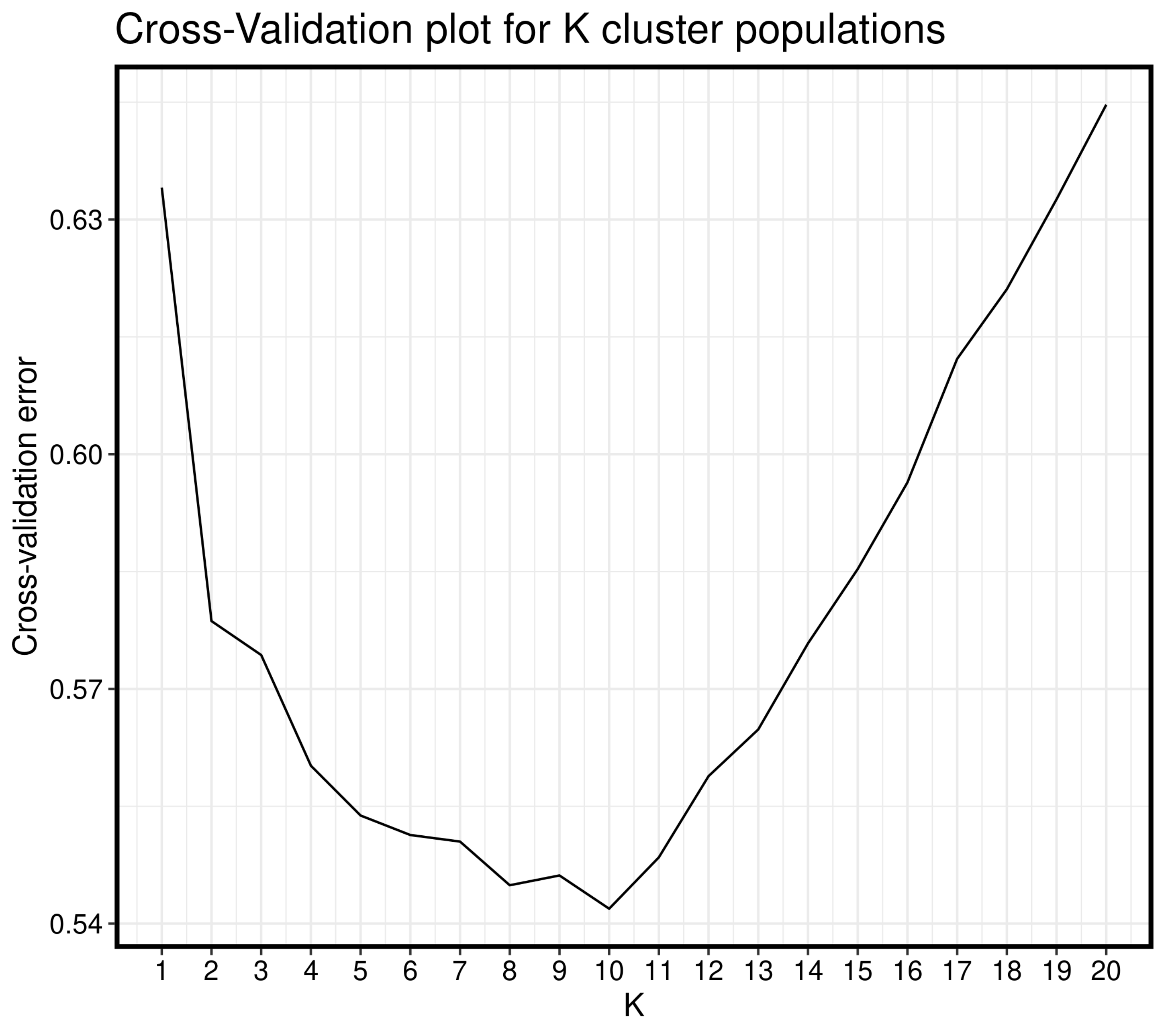
